# Characteristics of a novel NMR-based metabolomics platform for dogs

**DOI:** 10.1101/871285

**Authors:** Claudia Ottka, Katariina Vapalahti, Jenni Puurunen, Laura Vahtera, Hannes Lohi

**Affiliations:** PetBIOMICS Ltd, Helsinki, Finland; Department of Veterinary Biosciences and Department of Medical and Clinical Genetics, University of Helsinki, Helsinki, Finland; Folkhälsan Research Center, Helsinki, Finland

**Keywords:** canine, metabolomics, NMR, reference intervals, precision, stability

## Abstract

Metabolomics has proven itself an invaluable research tool, providing comprehensive insight to systemic metabolism. However, the lack of scalable and quantitative methods with known reference intervals and documented reproducibility has prevented the use of metabolomics in the clinical setting. This study describes the development and validation of a quantitative nuclear magnetic resonance (NMR) -based metabolomics platform for canine serum and plasma samples. Altogether 8247 canine samples were analyzed using a Bruker’s 500 MHz NMR spectrometer. Using statistical approaches derived from international guidelines, we defined reference intervals for 123 biomarkers, studied method precision, analyte storage stability, the effect of prolonged contact to red blood cells, differences of blood collection tubes, interference of lipemia, hemolysis and bilirubinemia, method comparison, and demonstrated the method’s practical relevance in a hyperglycemic cohort. Owing to the advantages of quantitative results, high reproducibility, and scalability, this canine metabolomics platform holds great potential for numerous clinical and research applications to improve canine health and well-being.

---

Metabolomics is an omics-based approach that generates comprehensive information on metabolism, enabling an extensive view on the current state of systemic metabolism. Metabolomics has become increasingly popular in canine studies. It is especially suitable in characterizing metabolic effects of multiple environmental and inter- and intra-individual^1^ factors including feeding^2–6^, aging^7^, inter-breed differences^8,9^, drug action^10,11^, behavior^12,13^, exercise^14^, genetic factors^15^ and pathological processes^16–26^.

Mass spectrometry (MS) and proton nuclear magnetic resonance (NMR) spectroscopy are the two main technologies for metabolic profiling, both in human and animal studies^27^. Proton NMR spectroscopy offers quantitative data with high reproducibility, high throughput and excellent scalability in a non-destructive and cost-effective manner^27^. NMR spectroscopy is well suited for the scientific use of large cohorts and biobanks^28,29^ and is highly suitable for studies combining different omics technologies^29–31^. Due to the quantitative and highly reproducible nature of NMR spectroscopy, it can be easily utilized as a diagnostic tool and research findings can be easily applied to clinical use.

Metabolomics holds great potential in clinical diagnostics. It is a promising tool for highlighting the metabolic alterations associated with the emergence and progression of diseases, early disease detection and individualization of treatment^32–34^. Clinical diagnostics and decision-making have formerly relied on the use of single clinical chemistry biomarkers. The routinely used approach is largely unusable for risk prediction, since changes are typically only seen when the diseased state has already been reached^33,35^. Thus, preventive and supportive care measures cannot be undertaken before organ failure is already present. Interpretation of the results by the traditional approach might also be challenging, since multiple simultaneous metabolic phenomena can affect the interpretation of these results and multiple diseases can be associated with similar laboratory results^22,35–37^. Moreover, evaluations of disease severity lack accuracy when only one affected metabolic pathway is taken into account^34,37^. Furthermore, traditional diagnostic approaches do not take the metabolic state of the patient into account, although it might have an impact on the best treatment of the patient^34,38^.

The prerequisites for the utility of a particular method include documentation of the method’s reproducibility, understanding of confounding factors that might affect the interpretation of the results, and the formation of reference intervals^39^. All of these factors have previously been largely unpublished in the canine metabolomics field. In this study, we describe the development and validation of a novel NMR-based metabolomics platform for dogs, demonstrate its performance characteristics, publish the determined dog-specific reference intervals for its biomarkers and demonstrate its utility in a practical situation.

## Results

To develop and validate a quantitative NMR-based approach for canine metabolomics, a total of 8247 serum or plasma samples were collected and analyzed. Thirty of these were pooled samples, 912 were replicate samples of client-owned dogs, 999 leftover clinical laboratory samples, and 6306 were single samples of client-owned dogs (Supplementary Figure 1). The number of individual client-owned dogs with samples collected during this project was 6164 (Supplementary Table 1). Eighteen percent of the dogs were under one year of age, 61% aged 1-7, and 21% over 7 years of age. Males consisted 40% of the population and samples were taken from 256 breeds.

### Proton NMR spectroscopy quantitates 123 biomarkers

All samples were analyzed with Bruker’s 500 MHz NMR spectrometer with spectral data interpreted by in-house scripts following principles demonstrated for human samples^40^. The NMR method was able to quantify 123 biomarkers, including extensive lipoprotein profiling, fatty acids, amino acids, albumin, creatinine and glycolysis related metabolites, in canine serum/plasma (Fig 1).

**Fig 1.**
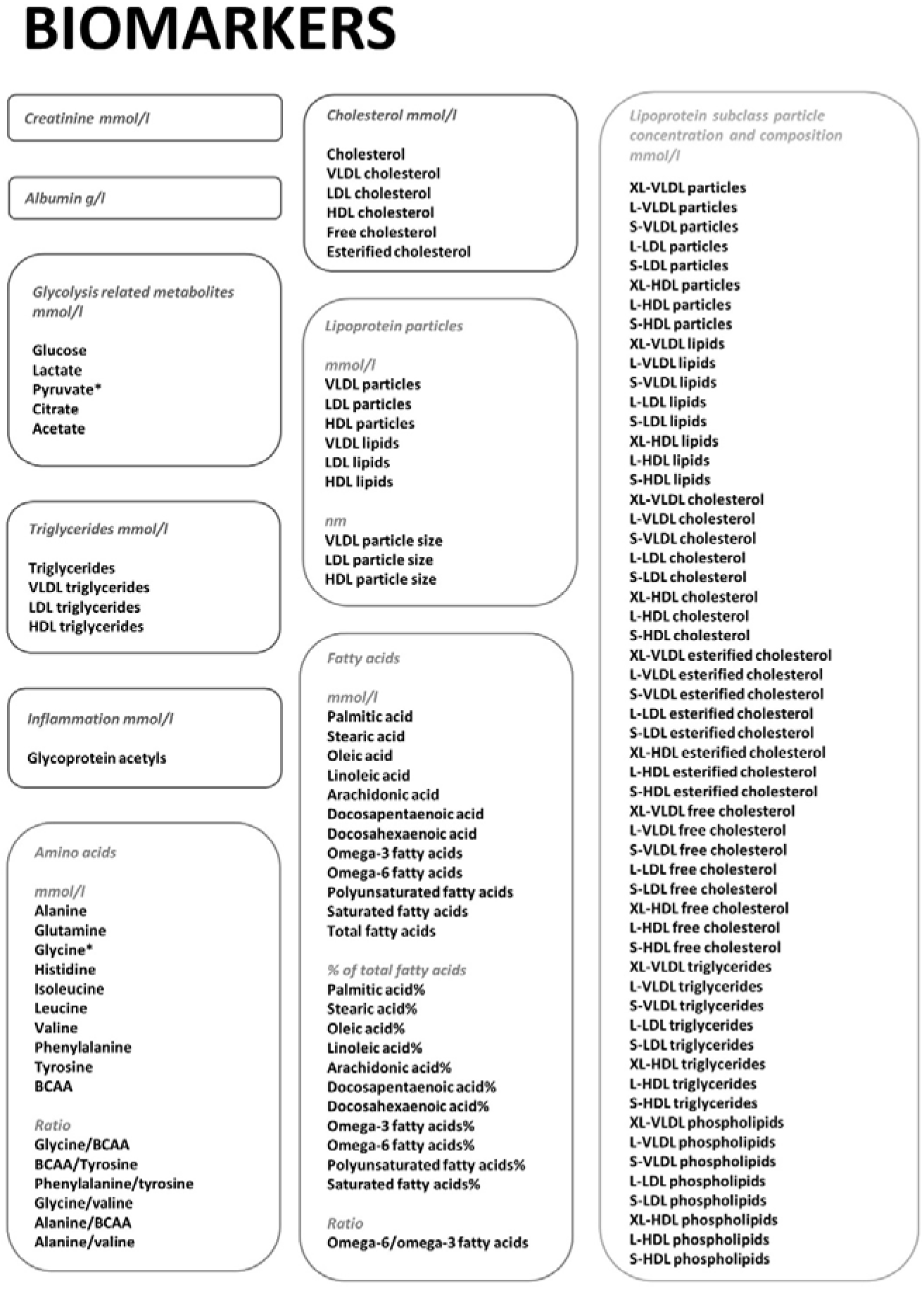
The biomarkers quantified by the NMR metabolomics testing platform. *: not available from EDTA-plasma samples. BCAA: Branched-chain amino acids XL-VLDL: Chylomicrons and very large VLDL-particles L-VLDL: large VLDL particles S-VLDL: small VLDL-particles L-LDL: large LDL particles S-LDL: small LDL particles XL-HDL: very large HDL particles L-HDL: large HDL particles S-HDL: small HDL particles

### Reference intervals for puppies, adult and senior dogs

We determined reference intervals (RI) (Table 1, Supplementary Table 2) with 90% confidence intervals (CI) (Supplementary Table 2) for 123 metabolites in serum and heparin plasma samples.

**Table 1.** Established reference intervals for the validated biomarkers.

| Analyte | HP all dogs | S all dogs | S puppy | S adult | S senior |
| --- | --- | --- | --- | --- | --- |
| Glucose mmol/l | 4.3-6.8 | 4.4-6.8 | 4.7-7.3<sup>a</sup> | 4.4-6.6 | 4.0-6.1 |
| Lactate mmol/l | 0.7-3.0 | 1.1-3.6 | 1.3-2.9<sup>b</sup> | 1.1-3.6 | 1.1-4.2 |
| Creatinine mmol/l | 40-99 | 32-103 | 21-96<sup>b</sup> | 40-108 | 37-104 |
| Albumin mg/dl | 26-32 | 25-32 | 22-31<sup>b</sup> | 26-32 | 26-32 |
| Cholesterol mmol/l | 3.8-10.4 | 3.6-10.3 | 3.8-10.1 | 3.6-10.3 | 3.5-10.6 |
| Triglycerides mmol/l | 0.22-0.97 | 0.19-1.00 | 0.18-0.76 | 0.19-0.97 | 0.19-1.13 |
| Pyruvate μmol/l | 29-155 | 11-107 | 9-96 | 11-111 | 13-109 |
| Citrate μmol/l | 63-122 | 61-123 | 57-115 | 61-124 | 62-128 |
| Acetate μmol/l | 19-36 | 21-37 | 22-40 | 21-37 | 20-37 |
| Alanine μmol/l | 214-584 | 216-597 | 205-504 | 205-583 | 244-650<sup>a</sup> |
| Glycine μmol/l | 147-466 | 130-454 | 145-594<sup>a,b</sup> | 127-381 | 128-403 |
| Glutamine μmol/l | 570-919 | 640-1,015 | 642-997<sup>b</sup> | 659-1,028 | 616-1,014 |
| Histidine μmol/l | 50-91 | 53-98 | 46-93 | 55-99 | 55-97 |
| Isoleucine $\mu\text{mol/l}$ | 33-80 | 37-89 | 30-80<sup>a</sup> | 38-92 | 38-87 |
| Leucine $\mu\text{mol/l}$ | 74-168 | 83-185 | 67-159<sup>b</sup> | 89-186 | 94-186 |
| Valine $\mu\text{mol/l}$ | 107-245 | 113-251 | 80-236<sup>b</sup> | 119-260 | 124-253 |
| Phenylalanine $\mu\text{mol/l}$ | 32-64 | 30-65 | 28-66 | 29-64 | 34-66 |
| Tyrosine $\mu\text{mol/l}$ | 39-85 | 41-89 | 39-83 | 41-90 | 46-93<sup>a</sup> |
| BCAA $\mu\text{mol/l}$ | 222-482 | 242-515 | 178-462<sup>b</sup> | 261-524 | 251-521 |
| Glycine/BCAA | 0.3-1.4 | 0.3-1.5 | 0.4-3.0<sup>a</sup> | 0.3-1.2 | 0.3-1.2 |
| BCAA/Tyrosine | 3.7-9.5 | 3.8-9.2 | 3.8-8.3<sup>a</sup> | 3.9-9.5 | 3.5-8.6 |
| Phenylalanine/tyrosine | 0.6-1.2 | 0.5-1.0 | 0.5-1.1 | 0.5-1.0 | 0.5-1.0 |
| Glycine/valine | 0.6-2.6 | 0.7-3.0 | 0.9-6.8<sup>a</sup> | 0.7-2.5 | 0.7-2.6 |
| Alanine/BCAA | 0.7-2.0 | 0.6-1.6 | 0.6-1.4 | 0.5-1.6 | 0.6-1.8<sup>a</sup> |
| Alanine/valine | 1.3-3.7 | 1.2-3.5 | 1.3-3.0 | 1.1-3.5 | 1.3-3.8 |
| VLDL lipids $\text{mmol/l}$ | 0.1-1.1 | 0.1-1.2 | 0.1-0.8 | 0.1-1.1 | 0.1-1.3 |
| VLDL particles $\text{nmol/l}$ | 15-54 | 12-54 | 14-45 | 12-51 | 12-67 |
| LDL lipids $\text{mmol/l}$ | 0.5-3.7 | 0.7-3.7 | 0.7-3.7 | 0.7-3.6 | 0.7-4.5 |
| LDL particles $\text{nmol/l}$ | 200-1,400 | 240-1,300 | 270-1,300 | 240-1,300 | 240-1,600 |
| HDL lipids $\text{mmol/l}$ | 7.7-14.6 | 6.9-15.1 | 7.2-15.0 | 6.9-15.1 | 6.6-15.3 |
| HDL particles $\text{nmol/l}$ | 32,000-57,000 | 30,000-58,000 | 33,000-58,000 | 30,000-59,000 | 28,000-58,000 |
| VLDL particle size $\text{nm}$ | 35.8-43.6 | 35.2-43.8 | 35.1-41.7<sup>a</sup> | 35.3-43.8 | 35.4-44.4 |
| LDL particle size $\text{nm}$ | 22.1-23.9 | 22.2-23.5 | 22.3-23.5 | 22.2-23.4 | 22.2-23.5 |
| HDL particle size $\text{nm}$ | 10.1-10.7 | 10.1-10.7 | 10.1-10.6 | 10.0-10.6 | 10.1-10.7 |
| VLDL cholesterol $\text{mmol/l}$ | 0.03-0.31 | 0.03-0.31 | 0.03-0.23 | 0.03-0.29 | 0.03-0.36 |
| LDL cholesterol $\text{mmol/l}$ | 0.16-2.35 | 0.28-2.27 | 0.30-2.24 | 0.28-2.21 | 0.28-2.85 |
| HDL cholesterol $\text{mmol/l}$ | 3.6-8.0 | 3.2-7.9 | 3.6-7.9 | 3.2-7.9 | 3.0-8.0 |
| Esterified cholesterol $\text{mmol/l}$ | 3.1-8.2 | 2.9-8.1 | 3.2-8.0 | 2.9-8.1 | 2.9-8.4 |
| Free cholesterol $\text{mmol/l}$ | 0.7-2.2 | 0.6-2.2 | 0.7-2.0 | 0.6-2.2 | 0.6-2.4 |
| VLDL triglycerides $\text{mmol/l}$ | 0.02-0.70 | 0.00-0.70 | 0.00-0.44 | 0.00-0.69 | 0.00-0.86 |
| LDL triglycerides $\text{mmol/l}$ | 0.13-0.30 | 0.13-0.31 | 0.15-0.33<sup>a</sup> | 0.13-0.29 | 0.14-0.27 |
| HDL triglycerides $\text{mmol/l}$ | 0.01-0.08 | 0.00-0.08 | 0.00-0.04<sup>b</sup> | 0.00-0.07 | 0.01-0.09<sup>a</sup> |
| Glycoprotein acetyls $\mu\text{mol/l}$ | 532-964 | 597-1,028 | 611-996 | 596-1,005 | 605-1,129<sup>a</sup> |
| Palmitic acid $\text{mmol/l}$ | 1.9-3.5 | 1.8-3.6 | 1.8-3.4 | 1.8-3.6 | 1.8-3.9 |
| Stearic acid $\text{mmol/l}$ | 1.9-3.8 | 1.7-3.8 | 1.8-3.7 | 1.7-3.9 | 1.7-4.0 |
| Oleic acid mmol/l | 1.1-2.5 | 1.3-2.8 | 1.3-2.5 | 1.3-2.8 | 1.3-3.2<sup>a</sup> |
| Linoleic acid mmol/l | 2.7-5.7 | 2.5-5.9 | 2.5-5.1 | 2.5-5.9 | 2.5-6.2 |
| Arachidonic acid mmol/l | 1.6-3.7 | 1.4-3.6 | 1.5-3.7 | 1.4-3.6 | 1.3-3.7 |
| Docosapentaenoic acid mmol/l | 0.1-0.3 | 0.1-0.4 | 0.1-0.3 | 0.1-0.4 | 0.1-0.4 |
| Docosahexaenoic acid mmol/l | 0.1-0.9 | 0.1-0.7 | 0.1-0.8 | 0.1-0.7 | 0.1-0.8 |
| Omega-3 fatty acids mmol/l | 0.4-1.8 | 0.4-1.6 | 0.4-1.6 | 0.4-1.6 | 0.5-1.8 |
| Omega-6 fatty acids mmol/l | 4.5-9.6 | 4.1-9.8 | 4.2-9.3 | 4.1-9.6 | 3.9-10.6 |
| Polyunsaturated fatty acids mmol/l | 5.2-10.8 | 4.7-11.1 | 4.8-10.6 | 4.7-11.0 | 4.6-12.0 |
| Saturated fatty acids mmol/l | 3.9-7.2 | 3.6-7.4 | 3.6-6.8 | 3.6-7.5 | 3.5-7.7 |
| Total fatty acids mmol/l | 10.5-20.4 | 9.7-21.0 | 9.8-20.0 | 9.7-21.2 | 9.3-22.5 |
| Palmitic acid% | 15.9-19.8 | 16-20 | 16-19 | 16-19 | 16-20 |
| Stearic acid% | 17.5-19.5 | 17.3-19.4 | 17.5-19.3 | 17.4-19.5 | 17.1-19.4 |
| Oleic acid% | 9.2-14.2 | 10.7-15.1 | 10.6-14.2<sup>a</sup> | 10.9-15.1 | 10.9-15.7 |
| Linoleic acid% | 23.9-29.4 | 24.1-28.6 | 23.3-27.5<sup>a,b</sup> | 24.5-28.6 | 24.7-28.7 |
| Arachidonic acid% | 13.6-20.7 | 13.2-20.1 | 15.0-20.7<sup>a,b</sup> | 13.1-19.9 | 12.7-19.2 |
| Docosapentaenoic acid% | 0.9-1.9 | 1.1-2.0 | 1.1-1.9 | 1.1-1.9 | 1.1-2.0 |
| Docosahexaenoic acid% | 0.9-6.2 | 0.7-4.8<sup>b</sup> | 1.0-5.2<sup>b</sup> | 0.7-4.7 | 0.6-5.0<sup>a</sup> |
| Omega-3 fatty acids% | 3-12 | 3-10 | 3-11 | 3-11 | 4-11 |
| Omega-6 fatty acids% | 43-49 | 42-48 | 42-48 | 42-48 | 42-47 |
| Polyunsaturated fatty acids% | 48-57 | 48-55 | 48-55<sup>b</sup> | 47-55 | 47-55 |
| Saturated fatty acids% | 34-39 | 34-38 | 34-38 | 34-38 | 33-38 |
| Omega-6/omega-3 fatty acids | 3.5-16.3 | 4.2-13.4 | 4.1-13.5 | 4.2-14.2 | 4.0-13.1 |
| XL-VLDL particles nmol/l | 0-2 | 0-1 | 0-1 | 0-1 | 0-2 |
| L-VLDL particles nmol/l | 1-15 | 0-16 | 0-11<sup>a</sup> | 0-15 | 0-19 |
| S-VLDL particles nmol/l | 11-38 | 10-39 | 13-34 | 10-37 | 10-47 |
| L-LDL particles nmol/l | 79-380 | 90-370 | 110-350 | 91-380 | 83-400 |
| S-LDL particles nmol/l | 89-1000 | 140-960 | 130-940 | 130-920 | 140-1,100 |
| XL-HDL particles nmol/l | 1-6,200 | 0-5,800 | 25-5,500 | 0-5,800 | 1-6,000 |
| L-HDL particles nmol/l | 21,000-34,000 | 20,000-34,000 | 20,000-34,000 | 19,000-35,000 | 18,000-34,000 |
| S-HDL particles nmol/l | 11,000-19,000 | 11,000-20,000 | 11,000-19,000 | 11,000-20,000 | 10,000-20,000 |
| XL-VLDL lipids mmol/l | 0-0.3 | 0-0.3 | 0-0.2 | 0-0.2 | 0-0.4 |
| L-VLDL lipids mmol/l | 0-0.6 | 0-0.6 | 0-0.4<sup>a</sup> | 0-0.6 | 0-0.7 |
| S-VLDL lipids mmol/l | 0.1-0.4 | 0.1-0.4 | 0.1-0.3 | 0.1-0.3 | 0.1-0.4<sup>a</sup> |
| L-LDL lipids mmol/l | 0.2-1.2 | 0.3-1.2 | 0.3-1.1 | 0.3-1.2 | 0.3-1.3 |
| S-LDL lipids mmol/l | 0.2-2.6 | 0.3-2.4 | 0.3-2.4 | 0.3-2.3 | 0.3-2.9 |
| XL-HDL lipids mmol/l | 0.6-5.2 | 0.6-5.0 | 0.6-4.7 | 0.6-5.0 | 0.7-5.2 |
| L-HDL lipids mmol/l | 5.4-8.1 | 5.0-8.3 | 5.0-8.1 | 4.9-8.3 | 4.8-8.3 |
| S-HDL lipids mmol/l | 1.3-2.1 | 1.3-2.2 | 1.3-2.1 | 1.3-2.2 | 1.1-2.2 |
| XL-VLDL cholesterol mmol/l | 0-0.06 | 0-0.06 | 0-0.04 | 0-0.06 | 0-0.08 |
| L-VLDL cholesterol mmol/l | 0-0.13 | 0-0.13 | 0-0.08 | 0-0.12 | 0-0.15 |
| S-VLDL cholesterol mmol/l | 0.01-0.14 | 0.02-0.15 | 0.02-0.11 | 0.02-0.14 | 0.02-0.18 |
| L-LDL cholesterol mmol/l | 0.03-0.70 | 0.07-0.68 | 0.11-0.61 | 0.07-0.68 | 0.07-0.76 |
| S-LDL cholesterol mmol/l | 0.09-1.64 | 0.16-1.53 | 0.16-1.50 | 0.16-1.48 | 0.18-1.88 |
| XL-HDL cholesterol mmol/l | 0.3-3.0 | 0.2-2.8 | 0.3-2.9 | 0.2-2.7 | 0.2-2.9 |
| L-HDL cholesterol mmol/l | 2.6-4.3 | 2.3-4.4 | 2.5-4.3 | 2.3-4.5 | 2.2-4.4 |
| S-HDL cholesterol mmol/l | 0.5-1.0 | 0.5-1.0 | 0.6-1.0 | 0.5-1.0 | 0.5-1.0 |
| XL-VLDL esterified cholesterol mmol/l | 0-0.03 | 0-0.03 | 0-0.02 | 0-0.03 | 0-0.03 |
| L-VLDL esterified cholesterol mmol/l | 0-0.05 | 0-0.06 | 0-0.04 | 0-0.05 | 0-0.07 |
| S-VLDL esterified cholesterol mmol/l | 0-0.09 | 0.01-0.09 | 0.01-0.08 | 0.01-0.09 | 0.01-0.11 |
| L-LDL esterified cholesterol mmol/l | 0.01-0.51 | 0.03-0.49 | 0.05-0.43 | 0.02-0.48 | 0.02-0.55 |
| S-LDL esterified cholesterol mmol/l | 0.05-1.14 | 0.10-1.09 | 0.10-1.06 | 0.10-1.06 | 0.12-1.35 |
| XL-HDL esterified cholesterol mmol/l | 0.2-2.3 | 0.2-2.1 | 0.2-2.1 | 0.2-2.1 | 0.2-2.3 |
| L-HDL esterified cholesterol mmol/l | 2.3-3.7 | 2.0-3.8 | 2.2-3.7 | 2.0-3.8 | 1.9-3.8 |
| S-HDL esterified cholesterol mmol/l | 0.4-0.8 | 0.4-0.8 | 0.5-0.8 | 0.4-0.8 | 0.4-0.8 |
| XL-VLDL free cholesterol mmol/l | 0-0.05 | 0-0.05 | 0-0.03 | 0-0.04 | 0-0.05 |
| L-VLDL free cholesterol mmol/l | 0-0.08 | 0-0.08 | 0-0.05<sup>a</sup> | 0-0.07 | 0-0.10 |
| S-VLDL free cholesterol mmol/l | 0.01-0.05 | 0.01-0.06 | 0.01-0.05 | 0.01-0.05 | 0.01-0.07 |
| L-LDL free cholesterol mmol/l | 0.03-0.20 | 0.04-0.20 | 0.05-0.18 | 0.04-0.20 | 0.04-0.21 |
| S-LDL free cholesterol mmol/l | 0.04-0.48 | 0.05-0.45 | 0.06-0.45 | 0.05-0.43 | 0.05-0.53 |
| XL-HDL free cholesterol mmol/l | 0.1-0.7 | 0.1-0.6 | 0.1-0.6 | 0.1-0.6 | 0.1-0.6 |
| L-HDL free cholesterol mmol/l | 0.3-0.6 | 0.3-0.6 | 0.3-0.6 | 0.3-0.6 | 0.3-0.6 |
| S-HDL free cholesterol mmol/l | 0.1-0.2 | 0.1-0.2 | 0.1-0.2 | 0.1-0.2 | 0.1-0.2 |
| XL-VLDL triglycerides mmol/l | 0-0.22 | 0-0.17 | 0-0.13 | 0-0.17 | 0-0.24 |
| L-VLDL triglycerides mmol/l | 0.01-0.41 | 0.01-0.42 | 0.00-0.27<sup>b</sup> | 0.01-0.39 | 0.01-0.49 |
| S-VLDL triglycerides mmol/l | 0.03-0.16 | 0.02-0.16 | 0.02-0.13 | 0.01-0.15 | 0.02-0.19 |
| L-LDL triglycerides mmol/l | 0.11-0.24 | 0.10-0.24 | 0.11-0.26<sup>a,b</sup> | 0.09-0.23 | 0.10-0.22 |
| S-LDL triglycerides mmol/l | 0.02-0.07 | 0.03-0.07 | 0.03-0.08 | 0.03-0.07 | 0.03-0.07 |
| XL-HDL triglycerides mmol/l | 0-0.02 | 0-0.02 | 0-0.02 | 0-0.02 | 0-0.03 |
| L-HDL triglycerides mmol/l | 0.01-0.03 | 0-0.03 | 0-0.02 | 0-0.03 | 0-0.04<sup>a</sup> |
| S-HDL triglycerides mmol/l | 0-0.04 | 0-0.03 | 0-0.02 | 0-0.03 | 0-0.04 |
| XL-VLDL phospholipids mmol/l | 0-0.05 | 0-0.05 | 0-0.04 | 0-0.05 | 0-0.09 |
| L-VLDL phospholipids mmol/l | 0-0.09 | 0-0.11 | 0-0.06 | 0-0.10 | 0-0.15 |
| S-VLDL phospholipids mmol/l | 0.01-0.07 | 0.01-0.08 | 0.01-0.06 | 0-0.08 | 0.01-0.10 |
| L-LDL phospholipids mmol/l | 0.08-0.33 | 0.08-0.32 | 0.09-0.33 | 0.07-0.30 | 0.08-0.35 |
| S-LDL phospholipids mmol/l | 0.08-0.87 | 0.12-0.84 | 0.12-0.80 | 0.12-0.80 | 0.12-1.01 |
| XL-HDL phospholipids mmol/l | 0.35-2.27 | 0.33-2.22 | 0.32-1.97 | 0.30-2.21 | 0.34-2.37 |
| L-HDL phospholipids mmol/l | 2.65-3.92 | 2.48-3.94 | 2.61-3.79 | 2.44-3.95 | 2.56-4.01 |
| S-HDL phospholipids mmol/l | 0.74-1.18 | 0.72-1.19 | 0.76-1.14<sup>a</sup> | 0.72-1.19 | 0.67-1.21 |

HP all dogs n = 269, S all dogs n =865, S puppy n =152, S adult n =545, S senior n =168.
HP: heparinized plasma
S: serum
BCAA: Branched-chain amino acids
<sup>a</sup>: no overlap between the 90% CI of the upper reference limit of this age group compared to adult dogs
<sup>b</sup>: no overlap between the 90% CI of the lower reference limit of this age group compared to adult dogs

In certain analytes, such as XL-VLDL variables, the concentration in healthy animals was very low, causing highly skewed distributions. Also, automatic rejection of extremely low values by the platform’s quality control, caused inability to calculate CIs with the used non-parametric method. For these analytes, the lower reference limit was rounded to 0. For a multitude of analytes, the 90% CI width was higher than 20% of the RI width, due to skewed or heavy tailed distributions. This was especially observed in RI for puppies and senior dogs, where the n count was lower (n < 170) than in adult dogs. However, for some analytes, it was also observed in RIs with very high n counts (n > 800).

RIs of puppies differed from the RIs of adult dogs in many analytes, such as glucose, lactate, creatinine, albumin, glycine, glutamine, leucine, valine, branched chain amino acids (BCAA), VLDL particle size, and HDL and LDL triglycerides. The higher reference limit (RL) of senior dogs exceeded the higher RL of adult dogs for alanine, tyrosine, alanine/BCAA, glycoprotein acetyls (GlycA), oleic acid, docosahexaenoic acid%, HDL triglycerides, S-VLDL lipids and L-HDL triglycerides.

### Precision of the platform

We studied analysis precision using three biologically different dogs, with duplicate aliquots of each dog’s sample analyzed once a day during a twenty-day period (Supplementary Table 3). The aliquots from a senior dog suffering from hyperadrenocorticism showed marked chylomicronemia, and was excluded from method precision estimations. Outlier removal slightly affected result interpretation, thus results were evaluated with outliers removed. In the primarily evaluated (non-chylomicronemic) samples, 110/123 biomarkers reached all of their precision goals. Albumin did not reach its primary precision goal of ASVCP CV_min_ based on biological variation, but reached all its other precision goals; CV% goal based on ASVCP Clinical Decision Limits and the S_T_ goal of under 1/8 reference width (S_Tmax_). Histidine, acetate, phenylalanine/tyrosine and BCAA/tyrosine met their primary precision goal of CV% under 20% (CV_maxBG_), but did not meet their secondary precision goal of S_Tmax_. 12/123 biomarkers did not reach their primary precision goal of CV_maxBG_, but reached the S_Tmax_ goal. All of these analytes were lipid analytes with high inter-individual biological variation and large reference interval width, and were in low concentrations in the tested samples. The CV_maxBG_ goal was thus regarded inappropriate for these analytes, since the variation in these analytes’ concentrations was not considered to affect clinical decision-making.

Chylomicronemia caused imprecision in multiple lipid measurements as well as in creatinine and the amino acids leucine and phenylalanine. Precision of glutamine was non-evaluable, since the olefin oligomer gel in the sample tubes inhibited correct glutamine estimation from the NMR signal. The precision of XL-VLDL phospholipids was non-evaluable due to a large amount of missing observations caused by physiologically very low analyte concentrations in healthy dogs.

### The effect of sample storage on biomarker stability

All evaluated analytes remained stable in refrigerated temperature for 7 days (Table 2, Supplementary Table 4). Most of the analytes remained also stable at room temperature for 7 days. Unstable metabolites in room temperature were the amino acids histidine, isoleucine, glutamine, tyrosine and phenylalanine, acetate, triglycerides and certain lipoprotein particle constituent concentrations as well as the fatty acid docosapentaenoic acid. The stability of all other amino acids than glutamine was better in EDTA-plasma than in serum. Glutamine concentration changed significantly at 4 days of storage at room temperature in EDTA-plasma samples and at 7 days of storage in serum samples. The acceptable change limit (ACL) of glutamine and XL-VLDL phospholipids could not be calculated using the analysis mean coefficient of variation (CVa), since these were not available from the precision study. The changes observed in triglyceride concentrations at room temperature originate from the computational quantitation based on lipoprotein particle distribution and the lipid quantity in the lipoprotein particle core and their outer surface. These are slightly affected by sample storage at room temperature, thus affecting triglyceride quantitation.

**Table 2.** Critical storage times for analytes sensitive to certain storage conditions in at least one sample type.

| Analyte | Temperature | Storage time<sub>Serum</sub> (p) | Storage time<sub>EDTA</sub> (p) | Storage time<sub>HP</sub> (p) |
| --- | --- | --- | --- | --- |
| Glutamine | RT | **7 days<sup>a</sup> (0.00<sup>\*</sup>)** | **4 days<sup>a</sup> (0.04<sup>\*</sup>)** | **4 days<sup>a</sup> (0.11)** |
| Histidine | RT | **3 days (0.00<sup>\*</sup>)** | NO | **3 days (0.03<sup>\*</sup>)** |
| Isoleucine | RT | **2 days (0.04<sup>\*</sup>)** | NO | 4 days (0.20) |
| Phenylalanine | RT | **3 days (0.00<sup>\*</sup>)** | NO | 7 days (0.06) |
| Tyrosine | RT | **3 days (0.01<sup>\*</sup>)** | NO | NO |
| HDL triglycerides | RT | **2 days (0.01<sup>\*</sup>)** | 4 days (0.07) | 4 days (0.20) |
| LDL triglycerides | RT | **2 days (0.01<sup>\*</sup>)** | **4 days (0.02<sup>\*</sup>)** | 2 days (0.20) |
| VLDL triglycerides | RT | **2 days (0.00<sup>\*</sup>)** | **2 days (0.04<sup>\*</sup>)** | 4 days (0.11) |
| Triglycerides | RT | **24 hours (0.04<sup>\*</sup>)** | **2 days (0.02<sup>\*</sup>)** | 2 days (0.20) |
| Acetate | RT | **4 days (0.02<sup>\*</sup>)** | **7 days (0.00<sup>\*</sup>)** | **4 days (0.03<sup>\*</sup>)** |
| Docosapentaenoic acid | RT | **7 days (0.03<sup>\*</sup>)** | NE | 7 days (0.20) |
| Citrate | -20°C | **1 week (0.00<sup>\*</sup>)** | NO | - |
| Acetate | -20°C | **3 months (0.00<sup>\*</sup>)<sup>b</sup>** | **6 months (0.00<sup>\*</sup>)** | - |
| Glycoprotein acetyls | -20°C | **1 month (0.00<sup>\*</sup>)** | **1 month (0.01<sup>\*</sup>)** | - |
| Omega-3 fatty acids | -20°C | **6 months (0.02<sup>\*</sup>)** | NE | - |
| HDL particle size | -20°C | **3 months (0.04<sup>\*</sup>)** | **3 months (0.03<sup>\*</sup>)** | - |
| HDL particles | -20°C | **6 months (0.00<sup>\*</sup>)** | **6 months (0.00<sup>\*</sup>)** | - |
| LDL lipids | -20°C | NE | **6 months (0.03<sup>\*</sup>)** | - |
| LDL particles | -20°C | NE | **6 months (0.03<sup>\*</sup>)** | - |
| LDL cholesterol | -20°C | NE | **6 months (0.03<sup>\*</sup>)** | - |
| VLDL particles | -20°C | **6 months (0.04<sup>\*</sup>)** | **6 months (0.01<sup>\*</sup>)** | - |

The storage time presented in the table represents the time point, where statistically significant changes (two-sided Wilcoxon exact test $p < 0.05$ ), with a mean percentage deviation (MPD) exceeding the acceptable change limit (ACL) are first observed. In sample types where statistical significance was not observed ( $p \geq 0.05$ ), the storage times represent the time point, where the MPD first exceeds the ACL. This table includes only primary analytes. Lipoprotein particle subclasses are not included. Stability at -20°C and -80°C were not studied in heparin plasma. $n_{\text{serum}} = 7$ , $n_{\text{EDTA}} = 7$ , $n_{\text{HP}} = 4$ .
RT: room temperature
\*: $p < 0.05$
NO: no change observed
NE: not evaluable due to discrepancies in p-values and MPD vs ACL. Results in other sample types should be consulted
until further studies are conducted.
<sup>a</sup> ACL unknown, evaluated only as $p < 0.05$ , time in heparin plasma represents the time, where a change in similar
magnitude as in serum and EDTA plasma is observed
<sup>b</sup> At 6 months of storage, MPD returned below the ACL and $p > 0.05$

All of the evaluated analytes were stable at −80°C for 12 months. Storage at −20°C affected certain lipoprotein particles and their composition, certain fatty acids, citrate, acetate and GlycA. Citrate levels were significantly (p < 0.05) changed already at one week of storage at −20°C.

Heparin plasma storage samples rarely made under the p-value of 0.05, even when the mean percentage deviation (MPD) exceeded the ACL and statistically significant (p < 0.05) changes were noted in other sample types.

Outlier removal slightly affected result interpretation in serum and EDTA plasma, thus their results were evaluated after outlier removal. In heparin plasma, outliers could not be reliably identified due to the small sample size (n = 4).

### The effect of delayed plasma separation on biomarker stability

We studied the effect of delayed plasma separation in 34 EDTA plasma samples, which had been stored as whole blood in the refrigerator for 24 and 48 hours before separating plasma (Supplementary Table 5, table 3). Outlier removal slightly affected result interpretation, thus the results were evaluated after outlier removal. Prolonged contact with red blood cells (RBC) affected the concentration of many of the analytes. Significant changes (p < 0.05, MPD>ACL) after one day’s contact to RBCs in refrigerator temperature were observed for glucose and lactate. After two days’ contact with RBCs, significant changes were observed for citrate, amino acids alanine, histidine, isoleucine, leucine, valine, phenylalanine and tyrosine, cholesterol, triglycerides, and lipoprotein particles and their constituents. Glutamine values were also changed significantly (p < 0.05) at 48 hours of storage as whole blood, but MPD could not be evaluated against the ACL due to the missing ACL.

**Table 3.** Critical storage times as whole blood for analytes sensitive to storage as whole blood.

| Analyte | Storage time (p) |
| --- | --- |
| Glucose | 24 hours (0.00) |
| Lactate | 24 hours (0.00) |
| Citrate | 48 hours (0.00) |
| Alanine | 48 hours (0.00) |
| Glutamine | 48 hours<sup>a</sup> (0.00) |
| Histidine | 48 hours (0.00) |
| Isoleucine | 48 hours (0.00) |
| Leucine | 48 hours (0.01) |
| Valine | 48 hours (0.00) |
| Phenylalanine | 48 hours (0.00) |
| Tyrosine | 48 hours (0.00) |
| HDL lipids | 48 hours (0.03) |
| HDL particles | 48 hours (0.00) |
| LDL diameter | 48 hours (0.00) |
| VLDL diameter | 48 hours (0.04) |
| Cholesterol | 48 hours (0.02) |
| HDL cholesterol | 48 hours (0.00) |
| Esterified cholesterol | 48 hours (0.00) |
| Triglycerides | 48 hours (0.00) |
| VLDL triglycerides | 48 hours (0.00) |

The storage time presented in the table represents the time point, where statistically significant changes (two-sided Wilcoxon exact test $p < 0.05$ ) together with a mean percentage deviation (MPD) exceeding the acceptable change limit (ACL) are first observed. $n = 34$ . This table includes only primary analytes. Lipoprotein particle subclasses are not included.
<sup>a</sup> ACL unknown

### Sample tube validation

We studied the differences of seven different sample tubes and the variability between their two sample lots using samples from 20 client-owned dogs (Supplementary Table 6). Outlier removal did not affect result interpretation, thus results were evaluated without outlier removal.

Firstly, the different sample tubes were compared to the primary reference tube (Vacuette Lithium Heparin). Most analytes showed comparable results in all tube types compared to the primary reference tube. Significant differences (p < 0.05) with MPD > ACL were noted for citrate, glucose, lactate and GlycA and significant differences (p < 0.05) with MPD < ACL were observed for glutamine, histidine, pyruvate and acetate. Glutamine showed significant differences (p < 0.05) compared to the primary reference tube, but its ACL was not available. Most of this variability was observed between analyte values in serum and plasma.

Secondly, analyte values in all serum tubes were compared to the reference serum tube (Vacuette Z Serum Clot Activator). All serum tubes showed comparable results with the serum reference tube for all metabolites.

The only lot-to-lot variability was observed in MiniCollect Serum gel tubes for glutamine and GlycA. Results obtained by both of these tube lots were still comparable to the reference serum tube (Vacuette Z Serum Clot Activator). In these MiniCollect Serum gel tubes, the two lots represented tubes of old-type (lot A) and new-type (lot B) tubes, with the tube having undergone changes in physical appearance, while sample tube constituents had remained the same.

### Interference studies

We studied the interference of hemolysis, lipemia, and unconjugated and conjugated bilirubin using triplicate samples of two different test pools for hemolysis and bilirubinemia and from one pool for lipemia (Supplementary Table 7). Hemolysis interfered with the quantitation of albumin and certain lipids; lipoprotein particle cholesterol concentrations, total triglycerides and certain lipoprotein triglyceride measures, multiple fatty acids, LDL and VLDL particle size, certain lipoprotein particle concentrations, lipoprotein phospholipids and certain lipoprotein lipids. Bilirubinemia interfered only with the quantitation of L-VLDL esterified cholesterol and S-HDL triglycerides. Lipemia affected the quantitation of GlycA due to the lipid contribution to the GlycA signal. However, the concentration of GlycA remained above the reference interval after the removal of lipids, indicating that lipemia was not the sole cause for raised GlycA concentration in the lipemic pool.

### Method comparison

Using 999 clinical canine samples, we evaluated, whether results of routine clinical chemistry analytes in the NMR metabolomics platform are comparable with results obtained by conventional clinical chemistry methods (Supplementary Table 8). The tested metabolites were glucose, lactate, creatinine, albumin, cholesterol and triglycerides. The tested samples covered the clinical concentration ranges for all analytes. The comparison plots showed satisfactory agreement between the two methods for all analytes. In some metabolites, logarithm transformation showed better agreement than initial data, suggesting better agreement in the transformed data.

Glucose, lactate and albumin measurements reached the Total Allowable Error (TE_a_) goals based on ASVCP Clinical Decision limits^41^, indicating that the two methods can be used interchangeably without affecting clinical decision-making. The bias percentage of creatinine, cholesterol and triglycerides was too high for the analytes to reach the TE_a_ goals based on Clinical Decision limits, meaning that the two methods cannot be used interchangeably. Values obtained by the conventional method can be evaluated against values obtained by the NMR method using the slope and intercept calculated by Deming regression.

### Hyperglycemia associates with many metabolic changes

We evaluated the practical relevance of the NMR metabolomics platform by comparing the metabolic profile of a markedly hyperglycemic group (n = 24) to a normoglycemic group of routine laboratory diagnostic samples (n = 781). Several biomarkers with previously reported changes in canine and human diabetes mellitus were significantly changed (p < 0.05) in the hyperglycemic group, including the BCAAs isoleucine, leucine and valine, GlycA, acetate, fatty acids, cholesterol, triglycerides, lipoprotein particle concentrations and composition, phenylalanine and lactate (Supplementary Table 9).

Excellent fit in logistic regression modeling (smallest AICs and AUC values > 0.9), indicating good predictive value for hyperglycemia, was observed for BCAAs and their ratios. Twenty metabolites had good fit (AUC < 0.8). 31 biomarkers had a high proportion of missing values, reducing the comparability with other metabolites due to different sample populations. There were 14 metabolites with good fit among these metabolites. Many of these metabolites are lipids, that have very low concentrations in healthy dogs, causing a high proportion of automatically rejected very low values by platform quality control.

## Discussion

Metabolomics is a rapidly growing field holding great potential for numerous clinical and scientific applications. Amongst various approaches, NMR spectroscopy is currently the most promising metabolomics method for clinical use due to its’ quantitative nature, high-throughput, accuracy and speed, all of which are currently not achievable with MS based metabolomics. In this study, we describe a novel, cost-effective NMR metabolomics testing platform for dogs. A similar approach has previously been demonstrated and widely utilized in human metabolomics^40^. The testing platform is capable of quantifying 123 biomarkers from a 100 μl sample of serum, heparin plasma, EDTA plasma or citrate plasma. The throughput for one device is ~200 samples per 24 hours and around 70,000 samples per year. The turnaround time is currently 5 days and can be markedly reduced. These characteristics make this comprehensive metabolomics platform a new high-throughput method to facilitate veterinary research and clinical diagnostics with significant implications on treatment, care and well-being of dogs.

Recent advances in metabolomics research are paving the way from diagnostics based on single biomarkers to a more holistic approach. Especially the management of chronic metabolic diseases is thought to benefit from a methodology capable of generating comprehensive information on the metabolic state of the individual, enabling a personalized approach to management of the disease^34^. Metabolomics is also proving itself valuable in disease risk prediction^33,42–44^. In contrast to genetics, metabolomics offers real-time information of the metabolic state of the animal, taking environmental factors and treatment into account. This offers metabolomics the possibility to be utilized as both a diagnostic tool as well as a follow-up tool. The advantage of the holistic nature of the NMR metabolomics platform was demonstrated in the comparison of hyperglycemic samples to normoglycemic samples from a routine laboratory diagnostic sample population. The NMR metabolomics platform was able to identify a plethora of biomarkers with significant differences between the hyperglycemic group and the normoglycemic group, including BCAAs, GlycA, acetate, fatty acids, cholesterol, triglycerides and lipoprotein particle concentrations and composition, phenylalanine and lactate. Many of these changes have been associated with canine and human diabetes mellitus, and even diabetes risk before disease onset^16,25,34,42,44^. This suggests the possibility of the NMR metabolomics platform to enhance preventative and precision veterinary medicine by providing new therapeutic targets and prognostic diagnostic markers.

In addition to a set of well-known clinical chemistry analytes, the established metabolomics platform includes a wide range of previously clinically unquantified biomarkers. One of the most interesting novel biomarkers is GlycA, a composite systemic inflammatory biomarker comprised of signals of acute-phase proteins α1 -acid glycoprotein, haptoglobin, α1-antitrypsin, α1-antichymotrypsin, and transferrin, with a slight contribution of glycosylated apolipoproteins^45^. In humans, GlycA has been linked with systemic inflammation, cardiovascular disease risk, diabetes mellitus, pregnancy, severe infection and all-cause mortality^28,45–49^. GlycA was also changed in our hyperglycemic group.

Formation of reference intervals is an important prerequisite for the clinical use of new laboratory method. We established reference intervals for 123 analytes, most of which are previously unpublished. The reference intervals differed between puppies, adult and senior dogs for many of the analytes, confirming substantial metabolic changes occurring during the maturation of the individual^50–53^. The higher reference limit of GlycA in senior than adult dogs suggests a higher prevalence of subclinical disease in older animals. The sample tube validation study demonstrated, that reference intervals created for the same sample type should be used in interpretation of the results. It should also be noted, that glycine and pyruvate cannot be quantified from EDTA plasma, and glutamine is not quantifiable from oligofin oligomer gel tubes.

Clinical use of a new laboratory method requires excellent method precision. The precision study suggested, that the precision of the NMR metabolomics platform is generally outstanding. For histidine, acetate, phenylalanine/tyrosine and BCAA/tyrosine, further studies in diseased animals are needed to conclude, whether measurement imprecision affects clinical utilization of these analytes.

The analytical method should be considered when evaluating laboratory results against treatment guidelines and reference intervals^39^. The method comparison study revealed a linear relationship between the conventional and NMR method for all analytes, but all results cannot be evaluated interchangeably. Triglyceride measurements showed most variability between the two methods, explained by differences in methodology, sample handling and biological characteristics. Analysis with the conventional method was done immediately, whereas NMR aliquots were frozen and one batch partly thawed during shipment. The methodology is also different, NMR measures triglycerides within lipoprotein particles, whereas the conventional method is based on interaction with the triglyceride molecule. Lipemia also affects the precision of triglyceride analyses, which causes more variability in high analyte concentrations.

Correct preanalytical measures are considered essential for the integrity of laboratory results^54,55^. An appropriate sample drawing technique is crucial for avoiding hemolysis, fasting before sample drawing for avoiding lipemia and prompt RBC separation for avoidance of prolonged RBC metabolism. Hemolysis, bilirubinemia, lipemia and prolonged contact with RBC all affected the quantification of certain metabolites. The profound impact of chylomicronemia on method precision was caused by matrix heterogeneity due to the chylomicron cream layer formation, which could be reduced by sample mixing immediately before analysis. Glycolysis-related metabolites are especially sensitive to contact with RBC, causing plasma samples to have different concentrations of these analytes than serum samples. Prolonged contact to RBCs affected also amino acids, which might explain why amino acid stability was better in our storage study using plasma and serum samples, than in previous studies using whole blood^56,57^.

It is critical to use sample storage and shipping conditions that preserve the integrity of laboratory results. All tested analytes were stable at refrigerator temperature for one week, making refrigerated temperature the optimal sample short-term storage and shipping temperature. Sample storage at room temperature should be minimized. Most amino were more stable in EDTA-plasma samples than in serum samples in room temperature, which has also been reported in previous studies^57^. The gold standard for long-term storage of serum/plasma metabolomics samples is freezing the samples immediately after serum/plasma separation and keeping them frozen at −80°C until analysis and avoiding additional freeze-thaw cycles^58^. All of the tested analytes remained stable during one year of storage at −80°C. Two weeks of storage in −20°C was suitable for most of the analytes.

Limitations of this study include, that the precision and interference studies did not include samples at clinical decision levels for all metabolites. Since the NMR platform is so extensive, acquiring these samples would have been extremely difficult. The sample numbers especially in heparin plasma, were a limitation of the storage study. Limitations of the method comparison study include lack of a reference method generating results that could be viewed as true values of the analyte. No clinical data was available for the hyperglycemia study, thus the study should be repeated with samples in confirmed diabetic animals. The identified analytes should also be studied for multicollinearity.

Owing to the advantages of quantitative results, high throughput and reproducibility, the NMR-based metabolomic platform developed and validated here holds great potential for numerous clinical and research applications in veterinary medicine. The performance of the NMR testing platform is generally outstanding and typical blood drawing and processing guidelines ensure the integrity of the results. The diagnostic power of the established metabolomics panel comes from the wide representation of markers from various molecular groups, including amino acids, fatty acids, glycolysis-related metabolites and lipoproteins. This enables efficient monitoring of physiologic changes in health and disease by the identification of condition-specific metabolic fingerprints.

## Materials and Methods

### Proton NMR spectroscopy

Metabolic profiling was conducted using a proton nuclear magnetic resonance (NMR) metabolomics technique similar to what has been demonstrated for human samples in Soininen et al.^40^. The fully automated process is capable of quantifying 200 samples per 24 hours and the throughput for one device is around 70,000 samples per year. Usable sample types are serum, EDTA-plasma, heparin plasma and citrate plasma. Since the concentration of the tested molecule is in a linear relationship with NMR signal intensity, the linearity of NMR results is considered inherently outstanding.

The NMR testing method was calibrated by firstly collecting 1008 canine EDTA-plasma samples and 120 serum samples from client-owned dogs and running the samples with NMR. Only samples with plasma/serum separation done within one day of sample collection were used for initial method calibration, resulting in 847 EDTA-plasma samples and all 120 serum samples. The characteristic NMR signals of amino acids, glycolysis related metabolites, creatinine, and GlycA are well-known, and the quantitative and linear relationship between the signal intensity and molecular concentration is an inherent property of NMR. Thus, these biomarkers can be quantified using the established NMR techniques. The identification of signals for triglycerides, cholesterol, and lipoprotein subclass particles and lipids was verified by analyzing 200 of the previously collected EDTA-plasma samples with different lipid concentrations by high-performance liquid chromatography^59^. For these measures, the concentration range of the automated NMR spectral analysis was also calibrated against these results.

Once the initial method calibration was completed, we collected a total of 4683 serum and 495 lithium heparin plasma samples across Finland during fall 2017-fall 2018 for use in our canine NMR metabolomics project. Samples were collected from dogs of all available breeds, with an emphasis of including dogs from genetically and morphologically different breed groups. Samples were drawn by cephalic venipuncture from client-owned dogs. Serum samples were collected into MiniCollect serum gel tubes and were allowed to clot for 30-45 minutes before centrifuging them at 3,000 x g for 10 minutes to separate the serum. Heparin plasma samples were collected into MiniCollect lithium heparin plasma gel tubes and the samples were immediately centrifuged at 3,000 x g for 10 minutes to separate plasma. All samples were then stored at −80°C before NMR analysis. All dog owners completed a history form, that included signalment and details about the health status, diet, exercise, stress and reproductive state of the dog.

The fatty acid panel verification and calibration was done using 100 samples with different lipid concentrations, collected for the canine NMR metabolomics project. Reference fatty acid concentrations were obtained using a chromatographical method by a commercial laboratory (Vitas as, Norway).

### Reference intervals

Determination of reference intervals (RI) of healthy dogs was performed according to the American College of Veterinary Clinical Pathology (ASVCP) reference interval guidelines^39^. We used the population-based nonparametric method for RI calculation, and calculated 90% confidence intervals (CI) for the reference limits. A minimum of 120 samples is required for the use of this method. Inclusion in RI calculation was based on fasting duration (minimum of 12 hours), appropriate sample handling, and the lack of owner-reported biological confounding factors, including diseases, severe anxiety/stress and strenuous exercise before blood collection.

A total of 865 samples collected for the canine NMR metabolomics project were included in the serum RI calculations. The samples were chosen to include over 120 samples of puppies (under 1 year old), adults (1-7 years old) and senior dogs (over 7 years old), to be able to calculate age-specific reference ranges. The 865 samples consisted of individuals from 68 breeds, 347 males and 517 females, 152 puppies, 545 adult dogs and 168 senior dogs.

A total of 269 samples out of the initial 495 samples collected for the canine NMR metabolomics project were qualified for lithium heparin plasma RI calculations. They consisted of individuals from 83 breeds, 155 males and 114 females, 29 puppies, 196 adult dogs and 44 senior dogs. Age-specific RI could be calculated for adult dogs.

Each metabolite was examined for outliers before RI calculation. We used Box Cox transformation to find a normal transformation, and Horn’s algorithm in the transformed data to identify the outliers. In Horn’s algorithm, the criterion for rejection of values is exceeding interquartile (IQ) fences set at Q1 - 1.5*IQR and Q3 + 1.5*IQR (IQR = interquartile range; IQR = IQ3 – IQ1 where IQ1 and IQ3 are the 25th and 75th percentiles, respectively. Once we identified the outliers, the sample and animal data were thoroughly reviewed for confounding analytical and preanalytical factors before reaching a conclusion about sample exclusion. In metabolites, in which normal distribution was not found, Horn’s algorithm offered multiple outliers, which all were reviewed before reaching a conclusion about exclusion. After calculating the RIs and the 90% CIs for lower and upper reference limits, we studied the relation between 90% CIs and RI. It is recommended that CI should not exceed 0.2 times the width of the RI, since it may indicate, that the sample number is insufficient.

Differences between RI for different age groups were evaluated by comparing the 90% CI of reference limits of puppies and senior dogs to the 90% CI of reference limits of adult dogs. Differences in the reference limits were concluded to be present, when there was no overlap between the 90% CI of reference limits between the two groups.

### Precision

Precision of the NMR method was tested using three biologically different dogs (puppy, healthy adult, senior dog suffering from hyperadrenocorticism). Blood was drawn by cephalic venipuncture into Vacuette 8ml Z Serum Sep Clot Activator tubes. Samples were allowed to clot for 30-45 minutes and centrifuged at 2,000 x g for 10 minutes to separate serum. Every sample was divided into 40 aliquots of 100μl. All samples were stored at −80°C before NMR analysis. Two duplicate aliquots of each sample were analyzed each day during a twenty-day time period.

The sample from the senior dog suffering from hyperadrenocorticism had marked chylomicronemia with a chylomicron coat forming on the top of the stored serum aliquots. The results of this sample were used to determine the effect of chylomicronemia on test precision.

Total within-laboratory precision estimates were expressed both as coefficient of variation (CV%) and standard deviation (S_T_). These within-laboratory precision estimates were evaluated against laboratory precision goals. We also calculated within-run precision, expressed as within-run standard deviation (S_r_). All precision estimates were calculated using methods described in CLSI EP5-A3^60^. Outlier detection was based on comparison of the absolute difference of the two duplicate aliquots analyzed in the same run to the mean absolute difference of all duplicates from the same sample. A difference that exceeded the mean absolute difference fourfold was set as the rejection line. If an outlier was detected, the duplicate was excluded from the calculations.

The currently most advisable hierarchy for performance specifications for human diagnostic laboratories is: 1. biological outcomes, 2. biological variation and 3. state-of-the-art^61,62^. Biological outcome-based performance goals cannot be used in veterinary medicine, since they have not yet been published for animals. Canine CV% goals based on biological variation are available for only a few analytes and can be so strict, that they are unachievable by clinical laboratories and do not affect interpretation of the results^41^. State-of-the-art goals can lack accuracy and might not fit all analytes. Due to the lack of specific performance goals for all analytes, we evaluated the precision results against a set of performance goals.

In this study, ASVCP goals based on biological variation were set as the primary goals for CV%, since they are the only canine-specific precision goals available^41^. The total accuracy of veterinary laboratory results is typically evaluated using Observed Total Error (TE_obs_) and evaluating it against against preset Total Allowable Error (TE_a_) limits. TE_obs_ can be calculated as TE_obs_ = Bias% + 2CV%. Bias was not applicable for this study as a deviation from a true value^41^. This led us to use ½ of ASVCP TE_a_ limits based on Clinical Decision Limits as our secondary precision goals for CV%. If ASVCP goals were not set, we used 20% as the goal for CV% (CV_maxBG_), which is generally considered an acceptable laboratory error in metabolomics^63,64^. Since individual CV% max goals were not available for most of the analytes and the 20% CV% goal might not be descriptive enough for metabolites that have a very wide or narrow reference range due to inter-individual biological variation, we additionally compared the S_T_ to the S_Tmax_ goal of 1/8 of the width of the reference range. This goal was based on the finding, that analytical variation over one quarter of the reference range is considered to affect the clinical interpretation of the results^65^. The S_Tmax_ goals based on 1/8 of the reference range width were very descriptive of the effects of imprecision on clinical interpretation of the results, but have the drawback, that imprecision affects the width of the reference range, itself^66^.

### Sample storage study

To define the effects of sample storage and shipping conditions on analyte results, we studied the effects of both long- and short-term sample storage. Short-term storage was studied in room temperature and refrigerated temperature, and long-term storage at −20°C and −80°C. To define storage effects in different sample matrices, we used three different sample matrices: serum, EDTA-plasma and heparin plasma. The protocol of the sample storage study is presented in Table 4.

**Table 4.** Sample storage study protocol.

| Storage time (T<sub>x</sub>) | Storage temperature |  |  |  |
| --- | --- | --- | --- | --- |
|  | Sample transport simulation |  | Sample storage simulation |  |
|  | Room temperature | Refrigerator | -20°C | -80°C |
| 0 hours (T<sub>0</sub>) | S, EDTA, HP |  |  |  |
| 8 hours | S, EDTA, HP | S, EDTA, HP |  |  |
| 24 hours | S, EDTA, HP | S, EDTA, HP |  |  |
| 48 hours | S, EDTA, HP | S, EDTA, HP |  |  |
| 72 hours | S, EDTA, HP | S, EDTA, HP |  |  |
| 96 hours | S, EDTA, HP | S, EDTA, HP |  |  |
| 1 week | S, EDTA, HP | S, EDTA, HP | S, EDTA | S, EDTA |
| 2 weeks |  |  | S, EDTA | S, EDTA |
| 1 month |  |  | S, EDTA | S, EDTA |
| 3 months |  |  | S, EDTA | S, EDTA |
| 6 months |  |  | S, EDTA<sup>a</sup> | S, EDTA |
| 12 months |  |  |  | S, EDTA<sup>a</sup> |

S: serum
EDTA: EDTA plasma
HP: lithium heparin plasma
<sup>a</sup> The EDTA plasma yield of two included dogs was slightly smaller than expected due to high hematocrit. For both dogs, one storage time point had to be excluded. The other sample was excluded from the 6 months storage of EDTA plasma at -20°C and the other one from 12 months storage of EDTA plasma in -80°C.

To study storage effects in these above-mentioned sample matrices, we took serum and EDTA plasma samples of seven client-owned dogs in a separate sampling situation, and heparin plasma samples of four dogs in another sampling situation.

During the sample collection of EDTA plasma and serum samples for the storage study, blood was drawn by cephalic venipuncture into Vacuette serum gel tubes and Vacuette EDTA K2 tubes. Samples were centrifuged and separated according to the tube manufacturers’ recommendations. EDTA-plasma samples were immediately centrifuged and plasma separated. Serum samples were allowed to clot for 30-45 minutes before centrifuging to separate serum. Separated plasma and serum were divided into 24 aliquots and stored in conditions outlined in Table 1 before immediate analysis with proton NMR spectroscopy.

Heparin plasma samples for the storage study were collected by drawing blood by cephalic venipuncture into Vacuette Lithium Heparin tubes. Samples were immediately centrifuged and plasma separated. The samples were divided into 13 aliquots and stored in conditions outlined in Table 4. After storage of samples in these test conditions, all samples were kept at −80°C for 2-3 weeks before NMR analysis.

We used the two-sided Wilcoxon exact test to estimate the statistical significance of the changes in analyte values during sample storage. The test was performed between samples not subjected to storage (T_0_) and samples subjected to different storage conditions at each storage time point (T_x_) in each storage temperature (Table 4). Two-sided p-values of Wilcoxon test statistic less than 0.05 indicated a statistically significant difference between metabolite values in T_0_ and T_x_.

In addition to the Wilcoxon test, we calculated the mean percentage deviation (MPD)

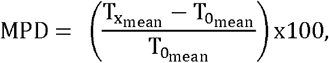

to indicate the stability/instability, as well as magnitude of change in metabolite values over a period of time^67^. MPD was compared to the acceptable change limit (ACL), according to ISO 5725-6^68^:

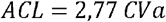

The CVa was obtained from the precision study results presented in this study, calculated as the mean CV% of the non-chylomicronemic samples. The factor 2,77 was derived from 1.96 x √2, where 1.96 represents the 95% of confidence interval for bi-directional changes, and √2 was used as we compared two results with the same CVa. A MPD higher than the ACL represents a probable change in analyte concentration.

Outliers were detected by duplicate analysis of samples in storage time T_x_ and collection time T_0_. Cut points to set the sample as an outlier were

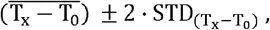

where 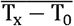 is the mean and 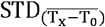 the standard deviation between samples in storage time T_x_ and collection time T_0_.

If the difference T_x_i__ – T_0_i__ oof the sample *i* exceeded or undercut these cut points, it was defined as an outlier. All analyses were done as outliers included or not excluded in purpose to study the effect of outlier exclusion.

The criteria for clinically significant change was set so, that the change should be both statistically significant (p-value of the two-sided Wilcoxon exact test <0.05) and the magnitude of the change, evaluated as MPD, was required to be above the ACL, and the change was required to be consistent at the remaining time points. Sample stability was studied only in primary analytes, analyte stability was not evaluated in ratio and percentage analytes.

### Delayed plasma separation study

In the delayed plasma separation study, we defined, how delays of plasma separation time from the sample tube manufacturer’s recommendations affect metabolite values. Delayed plasma separation was studied by storing samples as whole blood in refrigerated temperature for 24 and 48 hours before plasma separation.

Blood was drawn by cephalic venipuncture from 34 client-owned dogs into three Vacuette K2 EDTA tubes per dog. One of the tubes was centrifuged and plasma separated according to sample tube manufacturer’s recommendations within one hour after blood sampling. The remaining two tubes were stored as whole blood in a refrigerator until centrifugation and plasma separation after 24 and 48 hours of storage. After plasma separation, all samples were stored at −80°C before NMR analysis. This experiment was conducted using an older version of the NMR testing platform, thus the number of tested analytes was smaller, and fatty acids were not quantified.

For statistical analyses, we used similar methods as in the sample storage study. We used the two-sided Wilcoxon exact test to assess the effect of storage time on the sample, and mean percentage deviation (MPD) to indicate stability/instability and magnitude of change. Acceptable change limits (ACL) were the same as in the storage study. Outlier detection followed the same procedure as in the storage study, and criteria for clinically significant change were set similarly as p < 0.05 and MPD>ACL. T_0_ in the delayed plasma study represents plasma separated according to sample tube manufacturer’s recommendations and T_x_ plasma separated after 24 and 48 hours.

### Sample tube validation

To study, whether metabolite values differ between samples collected in different blood collection tubes (BCT), we studied both differences between different blood collection tubes and lot-to-lot variability within these tubes. The protocol was modified from the proposed blood collection tube validation process by Bowen and Adcock^69^.

Thirty milliliters of blood was drawn from a cephalic iv-cannula from 20 dogs into seven different types of blood collection tubes; MiniCollect Serum Separator Clot Activator tube, Vacuette Z Serum Separator Clot Activator tube, Vacuette Z Serum Clot Activator tube, Vacuette K2 EDTA tube, MiniCollect Lithium Heparin Separator tube, Vacuette Lithium Heparin tube and Vacuette Lithium Heparin Separator tube. Each tube type was tested with two tubes from different lots. EDTA and lithium heparin samples were immediately centrifuged to separate plasma. Serum samples were allowed to clot for 30-45 minutes before serum separation. The centrifugation conditions for samples collected into MiniCollect tubes were 3,000 x g for 10 minutes, and 2,000 x g for 13 minutes for all other sample tube types. The tested sample tubes and their specifications are presented in Table 5. All samples were stored at −80°C before analysis with NMR spectroscopy. The sample tube validation was only conducted regarding primary analytes, not for ratio and percentage analytes.

**Table 5.** Specifications of the tested blood collection tubes in the sample tube validation study.

| Blood Collection tube | Sample type | Clotting activator/anticoagulant | Gel type | Lots |
| --- | --- | --- | --- | --- |
| MiniCollect Serum Gel Tube 0,8ml | Serum | Clotting activator SiO<sub>2</sub> | Acrylic gel | A: 171010<br>B: A18023K3 |
| Vacurette Z Serum Separator Clot Activator 3,5ml | Serum | Clotting activator SiO<sub>2</sub> | Olefin oligomer gel | A: A17103HT<br>B: A18024EJ |
| Vacurette Z Serum Clot Activator tube 2ml | Serum | Clotting activator SiO<sub>2</sub> | - | A: A170837S<br>B: A180644P |
| Vacurette K2 EDTA Tube 2ml | EDTA-plasma | Spray-dried K2 EDTA 1,2-2 mg (waterless)/ 1 ml blood | - | A: A17063FX<br>B: A18043H3 |
| MiniCollect Plasma Gel Tube 0,8ml | Lithium heparin plasma | Spray-dried lithium heparin 18 I.U./1 ml blood | Acrylic gel | A: A17114HE<br>B: A18013N7 |
| Vacurette Lithium Heparin Tube 2ml | Lithium heparin plasma | Spray-dried lithium heparin 18 I.U./1 ml blood | - | A: A17083HR<br>B: A180247F |
| Vacurette Lithium Heparin Separator 3ml | Lithium heparin plasma | Spray-dried lithium heparin 18 I.U./1 ml blood | Olefin oligomer gel | A: A17123YY<br>B: A18053D8 |

We used the two-sided Wilcoxon exact test and mean percentage deviation (MPD) (detailed in 2.4 Sample storage study) to evaluate statistical significance of differences between the tubes.

Differences between different sample tubes were tested using this protocol:

1. Study, which tubes give comparable results with the primary reference tube The Vacuette Lithium Heparin tube was set as the primary reference tube (T_0_) and all other tubes (T_x_) were compared against it. Since we hypothesized, that most of the differences between sample tubes would originate from differences between serum and plasma, we secondly studied, which serum tubes give comparable results.
2. Study, which serum tubes give comparable results with the serum reference tube. The Vacuette Z Serum Clot Activator tube was set as the serum reference tube (T_0_) and all other serum tubes (T_x_) were compared against it.

The average of the analyte values in the two lots was used in these calculations.

Lot-to-lot variability was tested using this protocol:

1. Compare the two lots of each tube to another. Tubes in lot A were set as a reference tubes (T_0_) and tubes in lot B (T_x_) were compared against them
2. If lot-to-lot variability was observed, compare results of each lot to reference tubes. This was done to determine, whether samples from both lots give comparable results to the other tube types. Lots A and B were compared separately to the Vacuette Lithium Heparin and Vacuette Z Serum Clot Activator tubes.

The same outlier detection protocol was used as in the sample storage study (2.4), except that we used coefficient 3 for STD (instead of 2) in cut points: 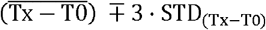. We set broader cut off limits for outlier detection for the sample tube validation than for the storage study, since the outlier test was more sensitive due to more observations in the sample tube validation than in the storage study.

In addition to the tests above, we calculated bias as mean difference of T_x_ and T_0_ with 95% confidence intervals, to highlight the limits in which the differences of the samples in tubes (or lots) T_x_ and T_0_ lie.

Although metabolite quantification is possible from citrate tubes, we did not include this tube type in this study, since this tube type is not generally used in clinical applications in this field.

### Interference

We tested our metabolites for interference of hemolysis, lipemia, and unconjugated and conjugated bilirubin. The testing protocol was derived from CLSI-EP7-A2^70^.

Hemolysis interference was studied using two client-owned dogs. 5 ml heparinized blood was collected by cephalic venipuncture for preparation of the hemolysis stock solution. The samples were centrifuged and plasma separated and discarded. The red blood cells were saved for the preparation of hemolysis stock solution. The hemolysis stock solution was prepared by the osmotic shock procedure. The hemoglobin concentrations of the hemolysates were measured by ADVIA 2120i hematology analyzer (Siemens)^71^. The base of the test and control samples were created by collecting 5ml heparinized blood by cephalic venipuncture from the same dogs that were used for the preparation of the hemoglobin stock solutions. Test samples containing 500 mg/dl hemoglobin were produced by adding a measured volume of the same dog’s hemolysate to the same dog’s plasma sample to produce the end concentration of 500mg/dl hemoglobin. Test samples containing 250mg/dl hemoglobin were prepared by mixing an equal amount of the same dog’s hemoglobin 500mg/dl test and control samples. All samples were divided into three aliquots and run as triplicates in the NMR analysis.

Bilirubin interference was studied by creating two sample pools (Pool 1 and Pool 2) from samples collected for the canine NMR metabolomics project. The sample pools were created so, that as many metabolite levels as possible would differ between the two pools. The baseline bilirubin concentrations of both pools were measured (540 nm) by a modified version of the acid diazo coupling (Malloy-Evelyn) method (Bilirubin Total (NBD), Thermo Fisher Scientific Oy, Finland), using a Konelab 60i chemistry analyzer^72^. The two pools were both divided into four pools, creating the bases for test and control pools for unconjugated bilirubin, and test and control pools for conjugated bilirubin. The test pools for conjugated bilirubin were prepared by adding bilirubin stock solution (201102 EMD Millipore) to the conjugated bilirubin test pool samples. The control pools for conjugated bilirubin were prepared by adding the same volume of water to the conjugated bilirubin control pool samples. The test pools for unconjugated bilirubin were prepared by adding bilirubin stock solution (2011 EMD Millipore) to the unconjugated bilirubin test pools. The control pools for unconjugated bilirubin were prepared by adding the same volume of NaOH to the unconjugated bilirubin control pool samples. All pools were divided into three aliquots and run as triplicates in the NMR analysis.

Lipemia interference was studied by pooling lipemic serum samples collected for the canine NMR metabolomics project. The test pool was created so, that the triglyceride concentration would be around 2mmol/l. The pool was divided into a test and control pool. The lipids of the control pool were cleared by ultracentrifugation. Both samples were divided into three aliquots and run as triplicates by NMR.

We used CLSI-EP7-A2^70^ guidelines as the base for our statistical testing. Interference testing was only evaluated regarding primary analytes, analyte stability was not evaluated in ratio and percentage analytes. Our testing was based on the point estimate of the observed interference effect (d_obs_) and the cut-off value (d_c_) and the comparison of these values.

The interference d_obs_ was defined as difference between the means of the test and control samples:

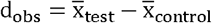, where test and control samples indicate samples with and without the interfering subject, respectively.

The cut-off value d_c_ was defined as 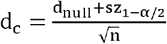, where d_null_ represented the assumed difference between the means of test and control samples (set to 0), n was the number of replicates (n=3) and s was the mean within-laboratory standard deviation (S_T_) of non-lipemic samples obtained from the precision study results presented in this study. 95% confidence intervals for the d_obs_ were calculated as

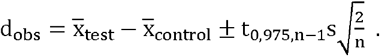

We established our decision of interference in the lower 95% limit of interference limit d_obs_low_ - interference was concluded to be present, if d_obs_low_ > d_c_ in at least one primary sample/sample pool, since properties of the sample matrix and/or analyte concentrations might affect the presence of interference. Hemolysis interference was concluded to be present if d_obs_low_ > d_c_ in at least the higher hemolysate concentration of 500mg/dl hemoglobin. Lipemia interference was only evaluated for molecules other than lipids, since the lipid removal process inevitably decreases the concentration of these molecules.

### Method comparison

The method comparison study was designed to evaluate, whether results from the routine clinical chemistry analytes included in the NMR metabolomics platform are comparable with results obtained by a conventional clinical chemistry method. The evaluated analytes were glucose, lactate, creatinine, albumin, cholesterol and triglycerides. In this study, bias was only evaluated as a measure of method interchangeability, since the values obtained by the conventional method cannot be regarded as true values of the analyte.

Clinical canine samples (n = 999) were studied by both NMR and standard clinical chemistry analysis methods. The samples included both heparin plasma and serum samples. The used samples were routine diagnostic sample material submitted by veterinarians across Finland, sent by mail to a single laboratory provider (Movet Ltd, Kuopio, Finland). Upon arrival, the samples were divided into two aliquots. One aliquot was immediately analyzed with conventional laboratory methods. Triglycerides were analyzed by a photometric method, cholesterol by a colorimetric method, albumin by a colorimetric method using bromcresol green, glucose by a photometric method using hexokinase, creatinine by a colorimetric enzymatic method and lactate by an enzymatic photometric method. The other aliquot was frozen and kept at −20°C for a maximum of 4 weeks before shipment to NMR analysis. The samples were sent in three batches on cool packs to NMR analysis, shipping time 7-14 hours. After arrival at the NMR laboratory, the samples were again stored at −80°C before NMR analysis. One of the sample shipments was sent over a 14h period on cool packs and partly thawed during shipment. Samples that were sent with a 7h shipment duration remained frozen during transportation. 404 of the samples were used for scaling the NMR albumin measurement, and thus excluded from the method comparison of albumin.

Statistical analysis followed the ASVCP General Quality Control Guidelines^73^. Agreement of the results was viewed using a comparison plot with values the NMR method plotted on the y-axis and the conventional method on the x-axis. Agreement plots were created for raw data and logarithm transformed data. Bland-Altman plots ^74^ were used to visualize the distribution of differences.

We utilized Deming regression^75,76^ in calculation of slope and intercept for all analytes, and in addition, linear regression for analytes with correlation over 0,99. Agreement of Deming transformed data was visualized by adding an agreement plot for data in which conventional method data was transformed using Deming regression coefficients. To determine the mean bias in the unit of the analyte at both ends of the reference intervals, bias was determined by slope and intercept of the Deming regression for these points:

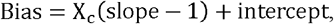

where X_c_ is the corresponding end of the reference interval.

To determine the interchangeability of results obtained by the conventional and NMR method, TE_obs_ was evaluated against ASVCP clinical chemistry guidelines based on Clinical Decision Limits^41^, which reflect the maximum values that would affect clinical decision-making. Observed Total Error TE_obs_ was calculated using the formula^41^:

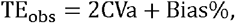

where CVa was the mean CV% of the non-lipemic samples in the precision study and the mean bias percentage (Bias%) was calculated using the formula ^41^:

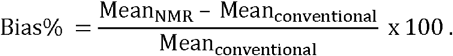

### Hyperglycemia-associated biomarkers

In order to demonstrate the utility of the developed method in an important biological question, we studied the differences in metabolite profiles in a hyperglycemic and normoglycemic group. We used the results of the 999 clinical canine samples utilized in the method comparison study as our base population for the study. No clinical data was available for the samples.

Diabetes mellitus is defined as a disorder, where blood glucose continuously rises over the renal glucose reabsorption threshold of 11mmol/l, causing glucosuria. The inclusion criteria to the case group (n = 24) was set as a NMR-measured glucose concentration over 11mmol/l. Inclusion to the control group (n=781) was based on the NMR-measured glucose being within the serum all dogs’ reference interval (4.4-6.8mmol/l). Since the control group consisted of clinical samples submitted to a clinical laboratory, the control group most likely includes animals suffering from diseases, that do not cause hyperglycemia.

To define metabolic changes associated with hyperglycemia, we calculated the means and their 95% CI for the case and control groups as well as the effect sizes of the observed changes. To visualize the differences in analyte concentrations between the two groups, we plotted histograms of the analyte data of the two groups. The significance of the difference between the metabolite values in the case and control group was studied using the two-sided Wilcoxon test’s Bonferroni-corrected p-values, with the cut-off for statistical significance set as p < 0.05.

To evaluate the metabolite’s ability of predicting hyperglycemia, we used single-variable logistic regression, in which case-control status served as the response variable and individual metabolites as dependent variables. Together with the metabolite’s maximum likelihood estimate’s p-value (significant < 0.05), area under the curve (AUC) values calculated for the model were used to determine the predictability of hyperglycemia, when the abnormal metabolite value is present. The higher the AUC value is (≤ 1), the better the fit of the model in terms of predicting ability; 0.7 being the cut point for fair, 0.8 for good and 0.9 for excellent model. Akaike information criterion (AIC) was performed in comparison of the models using different metabolites. It has no cut-off point and a lower AIC indicates better model, assuming that the test population in the models is the same. Glucose was excluded from this analysis, since it was used to define the case and control groups.

All calculations and plotting conducted throughout this study were performed by SAS version 9.4, SAS Institute Inc., Cary, NC, USA and Microsoft Office Excel, Microsoft Corp., Redmond, WA, US.

## Acknowledgements

We thank Kibble Labs Ltd. for the development and calibration of the canine NMR metabolomics platform, as well as NMR analysis of all samples. We thank Petra Jaakonsaari, Oona Mäki, Lea Mikkola, Marjo Hytönen and Maria Kaukonen for sample collection and Ileana Quintero for sample preparation for the interference experiment. Thomas Grönthal in the Veterinary Teaching Hospital, University of Helsinki, is acknowledged for performing the hemoglobin and bilirubin concentration measurements. We thank Sini Karjalainen, Kaisu Hiltunen, Reetta Hänninen, Juulia Tuominen, Kaisa Kyöstilä, Riika Sarviaho, Milla Salonen, and multiple veterinary students for help with sample collection and handling, and César Araujo for help with sample metadata management. We thank Tuomas Poskiparta, Katja Jauni, Jonas Donner, Heidi Anderson, Päivi Ruotanen, Julia Bouirmane, Essi Pekkala, Igor Polyakov, Leena Honkanen for help in participant acquisition, planning of data management and help in project conceptualization. We thank MediSapiens Ltd. for sample data and metadata management. We thank our collaborators Faunatar and ShowHau Center for enabling sample collection in their premises. We thank Movet Ltd. for the acquisition of the method comparison samples, and their analysis with conventional clinical chemistry methods. The authors also want to thank all dog owners, who offered their pets to participate in the study.

## Author Contributions

HL and CO conceptualized the study. CO, JP, LV, KV and HL designed the study. HL, CO, JP and LV participated in sample recruitment and collection. KV and CO conducted statistical analyses. HL supervised the study. CO drafted the manuscript with the help from other authors. All authors read and approved the final manuscript.

## Ethical Approval

All applicable international, national, and/or institutional guidelines for the care and use of animals were followed. All procedures performed in studies involving animals were in accordance with the ethical standards of the institution or practice at which the studies were conducted. Committee: Finnish national Animal Experiment Board, permit number: ESAVI/7482/04.10.07/2015.

## Data availability

The datasets generated during the current study are available from the corresponding authors upon request.

