## Supplementary Figure 1 for "Characteristics of a novel NMR-based metabolomics platform for dogs"

### Slide 1
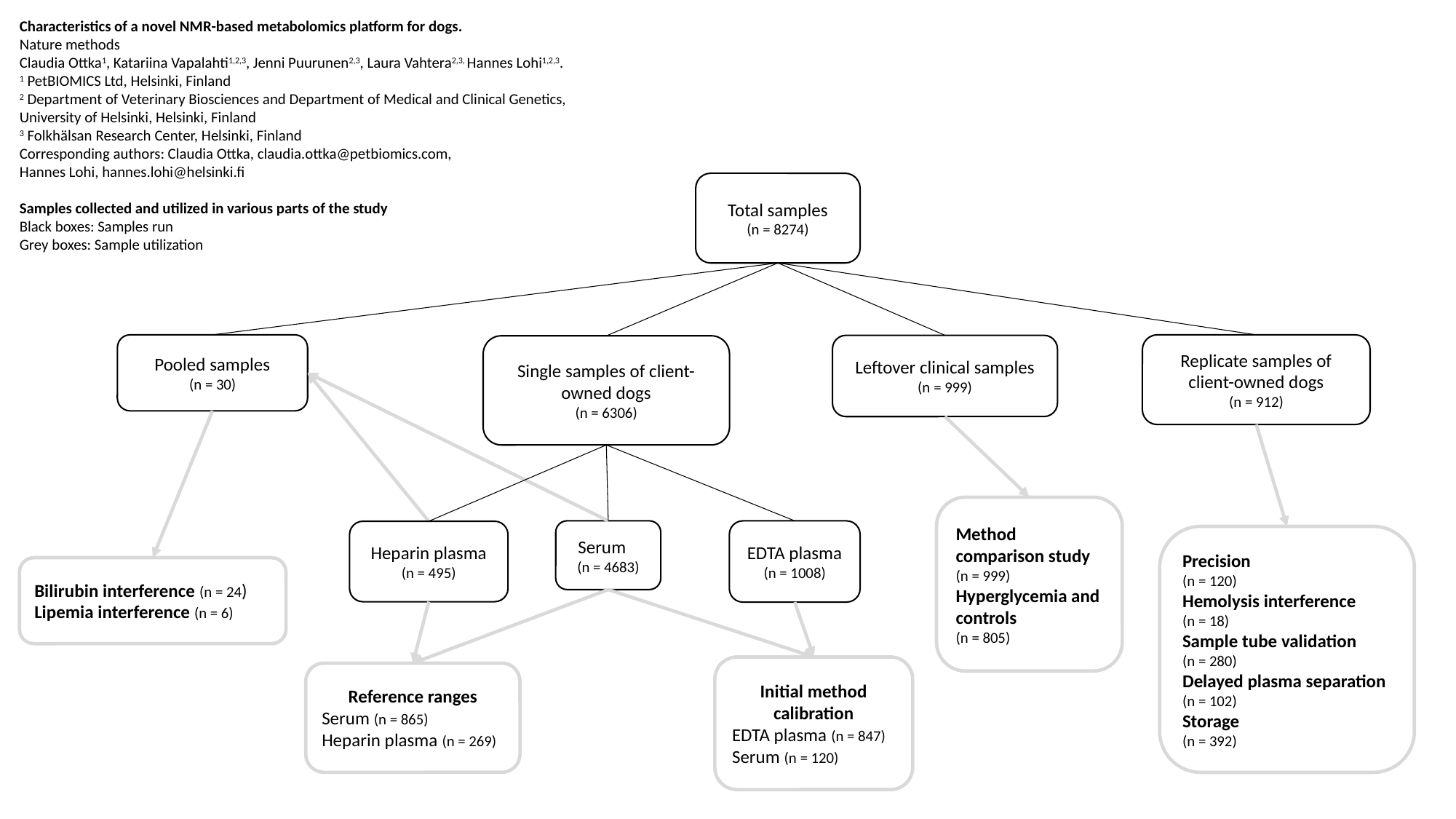

Characteristics of a novel NMR-based metabolomics platform for dogs.
Nature methods
Claudia Ottka^1^, Katariina Vapalahti^1,2,3^, Jenni Puurunen^2,3^, Laura Vahtera^2,3,^ Hannes Lohi^1,2,3^.
^1^ PetBIOMICS Ltd, Helsinki, Finland
^2^ Department of Veterinary Biosciences and Department of Medical and Clinical Genetics, University of Helsinki, Helsinki, Finland
^3^ Folkhälsan Research Center, Helsinki, Finland
Hannes Lohi,
Samples collected and utilized in various parts of the study
Black boxes: Samples run
Grey boxes: Sample utilization
Total samples
(n = 8274)
Pooled samples
(n = 30)
Replicate samples of client-owned dogs
(n = 912)
Leftover clinical samples
(n = 999)
Single samples of client-owned dogs
(n = 6306)
Method comparison study (n = 999)
Hyperglycemia and controls
(n = 805)
Serum (n = 4683)
EDTA plasma
(n = 1008)
Heparin plasma (n = 495)
Precision
(n = 120)
Hemolysis interference (n = 18)
Sample tube validation (n = 280)
Delayed plasma separation (n = 102)
Storage
(n = 392)
Bilirubin interference (n = 24)
Lipemia interference (n = 6)
Initial method calibration
EDTA plasma (n = 847)
Serum (n = 120)
Reference ranges
Serum (n = 865)
Heparin plasma (n = 269)
